# Targeting of the dosage-compensated male X-chromosome during early *Drosophila* development

**DOI:** 10.1101/671073

**Authors:** LE Rieder, WT Jordan, EN Larschan

## Abstract

The essential process of dosage compensation, which corrects for the imbalance in X-linked gene expression between XX females and XY males, represents a key model for how genes are targeted for coordinated regulation. However, the mechanism by which dosage compensation complexes identify the X-chromosome during early development remained unknown because of the difficulty of sexing embryos prior to zygotic transcription. We used meiotic drive to sex *Drosophila* embryos prior to zygotic transcription and ChIP-seq to measure dynamics of dosage compensation factor targeting. The *Drosophila* Male-Specific Lethal dosage compensation complex (MSLc) requires the ubiquitous zinc-finger protein Chromatin-Linked Adaptor for MSL Proteins (CLAMP) to identify the X-chromosome. We observe a multi-stage process in which MSLc first identifies CLAMP binding sites throughout the genome followed by concentration at the strongest X-linked MSLc sites. We provide insight into the dynamic mechanism by which a large transcription complex identifies its binding sites during early development.

## INTRODUCTION

Chromatin domains are enriched for specific histone modifications that activate or repress transcription, are a common property of metazoan genomes, and are critical for nuclear organization and regulation (Carelli et al., 2017). The formation of chromatin domains is often initiated early during embryogenesis (Evans et al., 2016; Vassetzky et al., 2000) However, little is understood about the dynamics of this essential process.

Strikingly-large chromatin domains include the dosage compensated chromosomes in heterogametic species, where all of the genes on a chromosome are coordinately regulated. For example, the human female inactive X chromosome is characterized by the heterochromatic marks H3K27 methylation and ubiquitinated H2AK119 (Hall and Lawrence, 2010). Like humans, *Drosophila melanogaster* also uses an X/Y system of sex determination, but males perform dosage compensation: male *Drosophila* upregulate their single X-chromosome approximately two-fold to equalize expression with that of females (Hamada et al., 2005; Larschan et al., 2011). The dosage compensated male X-chromosome is marked by acetylated H4K16 (Smith et al., 2001; Turner et al., 1992), which facilitates transcriptional elongation and increased expression (Zippo et al., 2009).

In *Drosophila*, Male-Specific Lethal complex (MSLc) is expressed only in males and accomplishes dosage compensation by specifically targeting the single X-chromosome. MSLc includes the histone acetyltransferase Males absent On the First (MOF) that deposits the activating H4K16ac mark (Akhtar and Becker, 2000; Smith et al., 2000). MSLc is most highly enriched at approximately 150-300 Chromatin Entry Sites (CES; also called High Affinity Sites) that contain GA-rich MSL Recognition Elements (MREs) (Straub et al. 2008; Villa et al. 2016; Alekseyenko et al. 2008). However, MREs are only two-fold enriched on the X-chromosome compared to autosomes (Kuzu et al., 2016). Furthermore, targeting of MSLc also requires the six-zinc finger transcription factor, Chromatin Linked Adaptor for MSL Proteins (CLAMP) (Soruco et al., 2013), although CLAMP targets MREs genome-wide and is not unique to the X-chromosome (Kaye et al., 2017; Rieder et al., 2017; Urban et al., 2017). Therefore, it remains unclear how MSLc targets the X-chromosome to establish an active chromatin domain.

Previous models for the formation of the dosage-compensated chromatin domain relied on steady-state patterns in mutant lines, often by examination of larval salivary gland polytene chromosomes. For example, while fully functional MSLc decorates the euchromatic region of the X-chromosome, partial complexes are recruited to a subset of X-linked CES (Deng et al., 2005; Gorman et al., 1995; Gu et al., 1998; Lyman et al., 1997; Palmer et al., 1994; Straub et al., 2008). CES are also the most highly enriched MSLc binding sites in wild-type flies (Alekseyenko et al., 2008). More recently, an *in vitro* DNA binding assay and the induction of MSLc in females both uncovered a subset of X-enriched sites called the Pioneering sites on the X (PionX) (Cheetham and Brand, 2018; Villa et al., 2016). Finally, when CES are synthetically inserted onto autosomes, they are targeted by MSLc, which then spreads in *cis* into neighboring chromatin as well as in *trans* to the male X-chromosome (Kelley et al., 1999; Larschan et al., 2007).

Based on these observations, several groups proposed a spreading model in which MSLc first targets CES or X-enriched sites and then spreads in two- or three-dimensions to active genes on the male X-chromosome (Alekseyenko et al., 2013; Lucchesi and Kuroda, 2015; McElroy et al., 2014; Ramírez et al., 2015; Soruco et al., 2013; Straub et al., 2008). However, directly testing this model *in vivo* is difficult: although dosage compensation is initiated early during development (Franke et al., 1996; Gergen, 1987; Polito et al., 1990; Rastelli et al., 1995), it is challenging to sex embryos pre-zygotic genome activation (ZGA).

To overcome this obstacle, we used a meiotic drive system to generate male and female-enriched pools of embryos and performed chromatin immunoprecipitation (ChIP)-seq for both CLAMP, MSLc and the H4K16ac chromatin mark at precise embryonic stages surrounding the initiation of dosage compensation. To our knowledge, this is the first time that sexing embryos prior to ZGA has been used to measure the recruitment dynamics of a dosage compensation complex in any species. We identified the following multi-stage process for targeting of MSLc: 1) CLAMP targets loci genome-wide; 2) MSLc identifies loci that are bound by CLAMP genome-wide, but not at CES; 3) MSLc becomes more enriched at CES and less enriched at other CLAMP binding sites. Overall, we provide insight into the dynamic mechanism by which a large transcription complex identifies its binding sites during early development.

## RESULTS AND DISCUSSION

### A meiotic drive system generates sexed embryos prior to ZGA

A significant impediment to observing the initial steps of dosage compensation is the inability to sex the *Drosophila* embryo before ZGA when sex-specific reporter transgenes are activated (Schauer et al., 2017). To overcome this obstacle, we used the Segregation Distorter (SD) meiotic drive system (Larracuente and Presgraves, 2012). In this system, expression of the *segregation distorter* gene product (Sd), a mislocalized form of RanGAP, interacts with sensitive alleles of the *Responder* (*Rsp*) locus, a large pericentromeric stretch of satellite DNA. In its naturally occurring form, both the *Sd* and *Rsp* loci reside on Chromosome 2, and Sd selfishly enriches its own transmission by preventing the maturation of spermatids carrying *Rsp*-bearing chromosomes.

Although the mechanism of SD remains unknown, the *Sd* and *Rsp* loci can be placed on ectopic chromosomes with the same effect. We therefore used existing *D. melanogaster* stocks in which *Sd* remains on Chromosome 2, but sensitive alleles of the *Rsp* locus (*Rsp^S^*) now reside on the X- or Y-chromosomes (Cheng et al., 2016; Polito et al., 1990; Walker et al., 1989). Males produced from this crossing strategy (**Figure 1A**) produce predominantly X- or Y-chromosome bearing sperm, depending on their genotypes, and sire a preponderance of progeny of the corresponding sex (**Figure 1B**). It should be noted that the data presented in **Figure 1B** represent our most conservative estimate of sex enrichment, and that we routinely observed much stronger drive.

**Figure 1.**
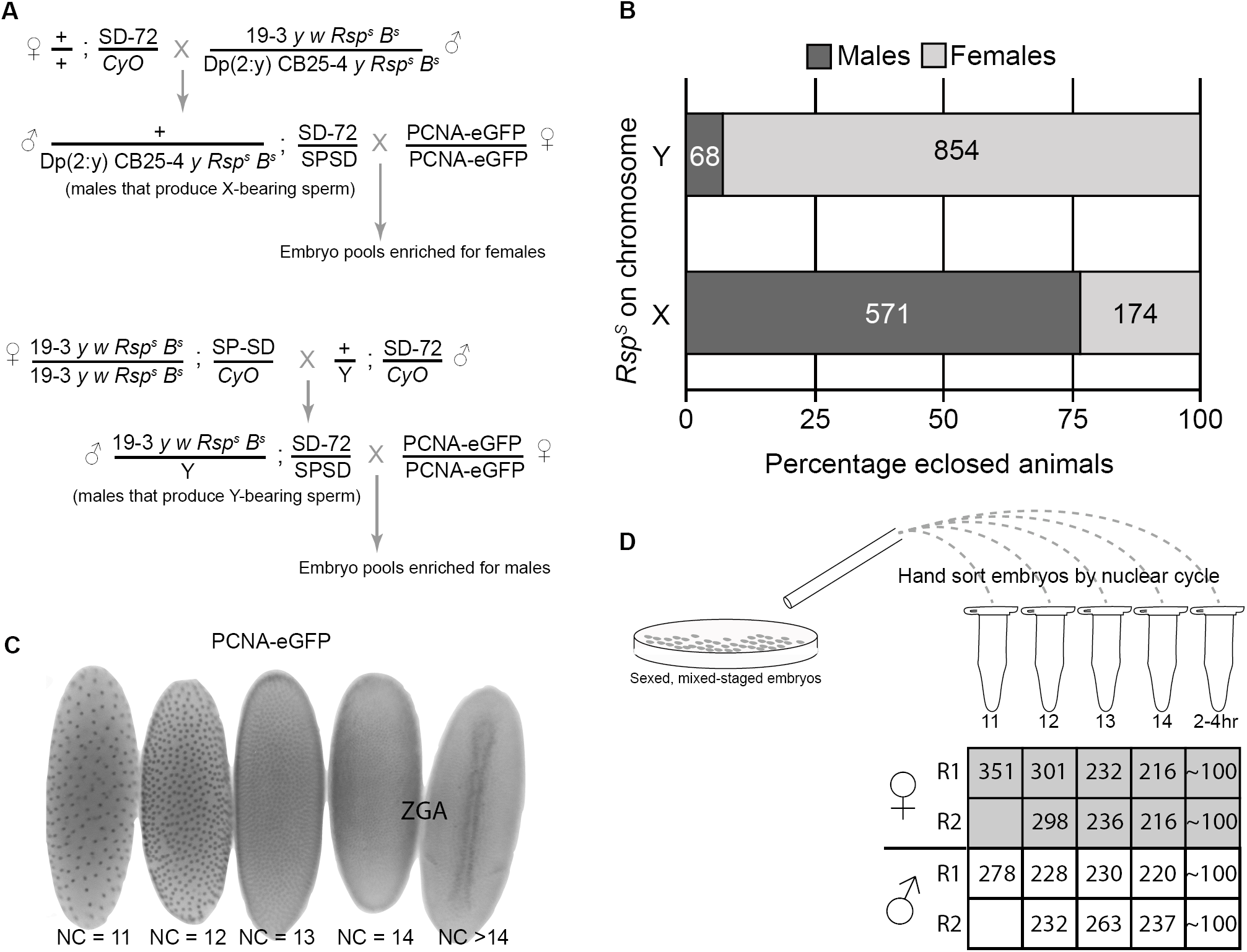
Validation of a Meiotic Drive System to Generate Sexed Embryos. (A) Crossing scheme using SD meiotic drive system to generate sex-enriched pools of embryos expressing PCNA-eGFP. *Rsps* is a sensitive allele of the *Rsp* locus. SPSD and SD-72 are strong Sd alleles and we found the drive to be strongest when the SD alleles were in *trans*. (B) Percentage of adult animals eclosed of each sex due to crossing schemes in (A). Numbers within bars represent total number of animals counted. (C) Staging embryos by nuclear cycle (NC) using the PCNA-eGFP transgene. Zygotic genome activation (ZGA) occurs during NC14. (D) Workflow to sort embryos for small-scale ChIP-seq. We hand sorted between 200 and 351 embryos for each biological replicate. We performed 2 biological replicates for male and female embryos between NC12 and 14 and one replicate for NC11. To represent a “late” stage of dosage compensation, we chose ∼100 2-4 hour embryos.

We combined the SD sexing system with a maternally-inherited PCNA-eGFP reporter transgene (Blythe and Wieschaus, 2016), which allowed us to precisely stage embryos by nuclear cycle, representing a developmental time course of approximately one hour surrounding ZGA (**Figure 1C**). These data demonstrate that, when combined, the SD meiotic drive and PCNA-eGFP reporter systems represent a powerful tool to both stage and sex embryos prior to ZGA. We hand sorted fixed embryos produced from these crosses (**Figure 1A**) and performed low input ChIP-seq (**Figure 1D**) (Blythe and Wieschaus, 2015) for factors associated with dosage compensation.

### MSLc identifies the male X-chromosome in a multi-stage process

We previously demonstrated that the CLAMP protein, which targets CES and other GA-rich sequences genome-wide, facilitates MSLc recruitment and dosage compensation (Kaye et al., 2018; Kuzu et al., 2016; Soruco et al., 2013). In addition, CLAMP is maternally deposited into the oocyte, while MSLc does not assemble until later in development (Graveley et al., 2011). We therefore hypothesized that CLAMP identifies its binding sites prior to MSLc. To test this hypothesis, we performed ChIP-seq on male-enriched embryo pools for CLAMP, MSL3 (a component of MSLc), and the H4K16ac mark, and on female-enriched embryo pools for CLAMP (Rieder et al., 2017) and H4K16ac. MSLc does not assemble in female embryos due to post-transcriptional repression of the structural MSL2 component by the master sex regulator Sex lethal (Kelley et al., 1997).

To perform small-scale ChIP-seq, we used 200-351 embryos for each nuclear cycle (**Figure 1D**). In addition, we performed ChIP-seq on approximately 100 sexed, mixed-stage embryos aged to 2-4 hours after egg lay, representing a post-ZGA stage (Gergen, 1987). Because hand sorting is laborious, we were limited in the number of biological replicates that we could perform: we performed two biological replicates for each sex and time point, except for nuclear cycle 11 and the 2-4 hour timepoint, for which we performed one biological replicate for each sex. In total, we hand sorted over 3,500 embryos. To ensure that we used only the highest quality data, we selected the best biological replicate based on which replicate had the highest overall scores for PCR bottlenecking coefficients 1 and 2 (measures of approximate library complexity) (Landt et al. 2012; Bailey et al. 2013) (**Table S1**). Overlap between peaks in selected samples is shown in **Figure S1**.

To visualize dynamic recruitment throughout early development, we plotted heatmaps of the highest quality replicates to measure occupancy at the following classes of loci: 1) PionX sites (Villa et al., 2016); 2) CES (Soruco et al., 2013); 3) CLAMP binding sites at the onset of ZGA identified in male nuclear cycle (NC) 14 (this study). We observed MSLc targeting in multiple stages. As we hypothesized, CLAMP identifies binding sites in both sexes prior to MSLc (**Figure 2A, B**). Before MSLc identifies CES in NC14, CLAMP is de-enriched at CES compared to other CLAMP binding sites. Concurrently, MSLc identifies other CLAMP binding sites genome-wide and is not restricted to the X-chromosome (**Figures 2C, S2**), an observation supported by male H4K16ac ChIP (**Figure 2D**). Because H4K16ac can be deposited by a different transcription complex in females (Non-Specific Lethal complex) (Raja et al. 2010), its occupancy pattern differs between males and females (**Figure 2D, E**) and is consistent with previous reports in cultured female Kc cells (Gelbart et al. 2009).

**Figure 2.**
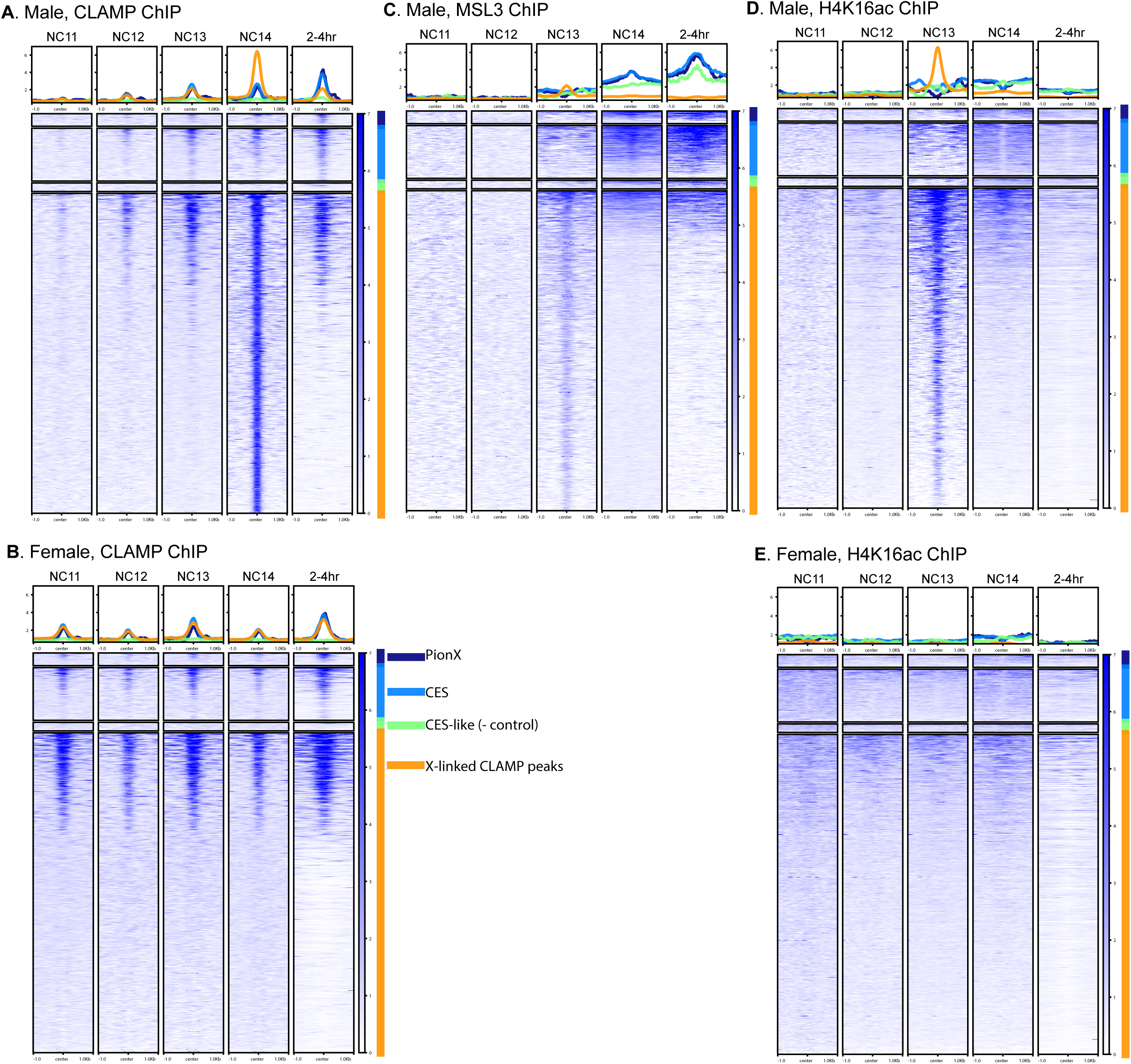
Staged, sexed ChIP-seq heat maps at X-linked CLAMP sites. Data are mapped over X-linked CLAMP sites and broken into categories, including PionX sites (Villa et al. 2016) (N=55; dark blue), CES (Soruco et al. 2013; Alekseyenko et al. 2008) (N=234; light blue), and other X-linked CLAMP peaks from the male NC14 sample (N=1417; orange). CES-like sites represent negative control sites (Soruco et al. 2013; Alekseyenko et al. 2008) (N=38; green). (A) CLAMP ChIP-seq from male embryos. (B) CLAMP ChIP-seq from female embryos. (C) MSL3 ChIP-seq from male embryos. (D) H4K16ac ChIP-seq from male embryos. (E) H4K16ac ChIP-seq from female embryos.

After MSLc identifies CES, CLAMP also becomes enriched at CES, specifically in males, consistent with the documented synergy between the two factors (Albig et al., 2019; Larschan et al., 2012). Individual profiles surrounding selected CES (**Figure S3**) are consistent with heat maps. Overall, we observe a dynamic pattern of MSLc targeting in which it first identifies CLAMP binding sites throughout the genome followed by specific targeting to CES. We do not observe a time point at which PionX sites (Villa et al. 2016) are specifically enriched for MSLc, compared to other CES.

In order to define how CLAMP, MSLc, and H4K16ac occupancies change over time at both genes on the X-chromosome and autosomes, we generated average gene profiles. CLAMP is similarly enriched on the X-chromosome (**Figure 3A, B**) and autosomes (**Figure S2A, B**) in both males and females, although CLAMP becomes enriched at CES compared to other binding sites after 2 hours (**Figure 2A, B**). CLAMP average gene profiles are similar in males and females until NC14 when the profiles diverge (**Figure 3A, B**), consistent with previously-documented synergy between CLAMP and MSLc only in males (Soruco et al. 2013). In contrast, MSLc becomes more enriched at CES and less enriched at other CLAMP binding sites over time (**Figures 2C, 3C**). MSLc and H4K16ac (**Figure 3D**) are enriched over gene bodies in males, consistent with previous reports in cell culture (Alekseyenko et al., 2006). Furthermore, the X-enrichment of MSLc and H4K16ac increases over time in males but not in females (**Figures 2C, D, 3C, D**).

**Figure 3.**
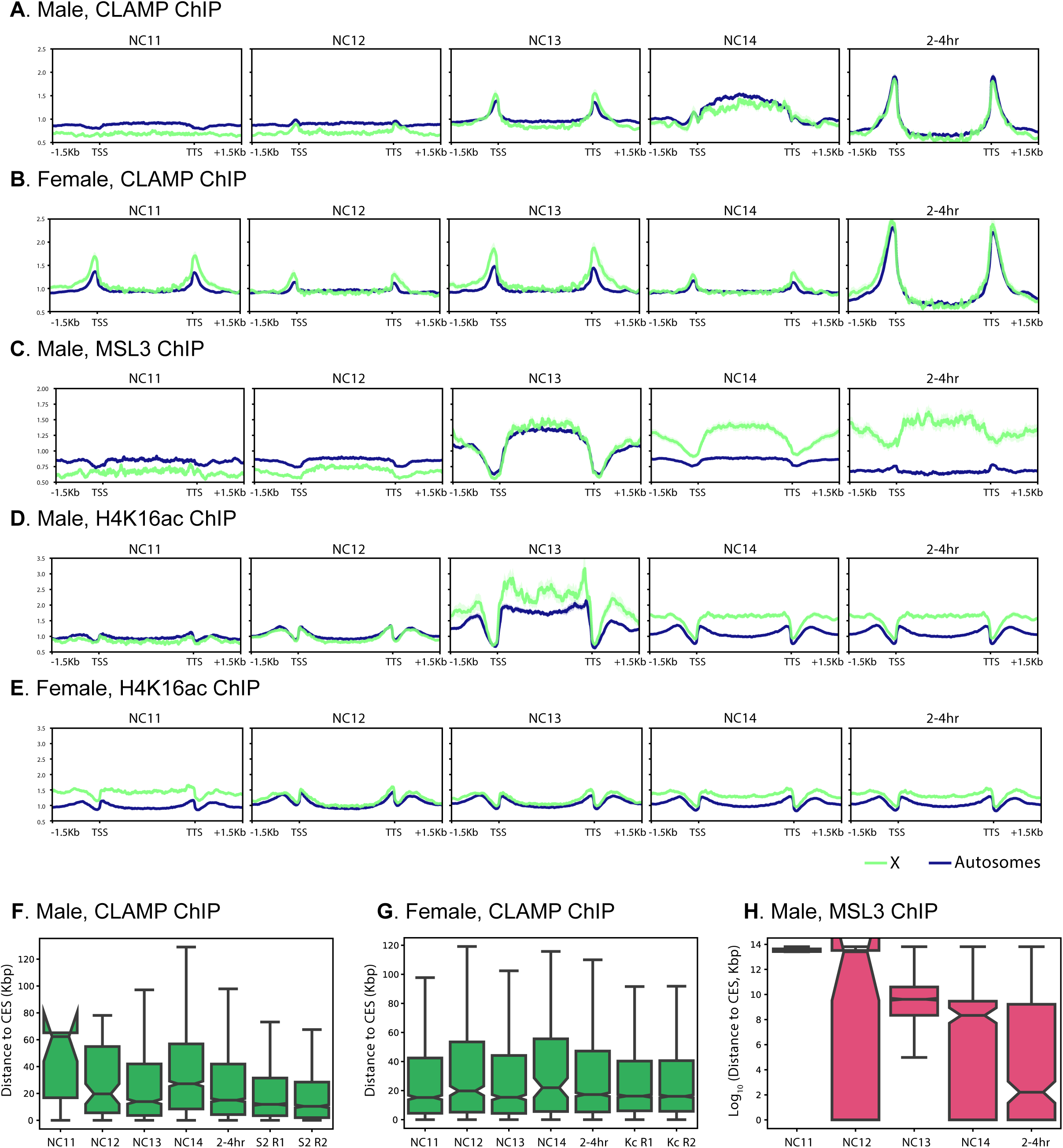
Average X-linked (green) and autosome-linked (blue) gene profiles over developmental time (A-E). TSS, transcription start site; TTS, transcription termination site. Distance to CES of (F) male CLAMP peaks, (G) female CLAMP peaks. For comparison, we have included previously published CLAMP ChIP-seq data from cultured male S2 cells and female Kc cells (two biological replicates each from Soruco et al. 2013). (H) Distance to CES of male MSL3 peaks.

We also investigated the distances of CLAMP and MSL3 peaks to the nearest CES over developmental time. In both males and females, CLAMP peaks reside very close to CES, even at the earliest time points (**Figure 3F, G**). However, male MSL3 peaks grow closer to CES throughout development (**Figure 3H**). Together these data suggest that CLAMP targets CES in both sexes early during development before MSLc complex identifies CES. Initially, MSLc identifies CLAMP binding sites genome-wide that are not located at CES. Next, synergy between CLAMP and MSLc occurs specifically at CES, and likely other factors, such as 1.688 repeat elements (Joshi and Meller 2017), enrich MSLc occupancy specifically at CES.

Overall, we define a multi-step process by which MSLc first identifies thousands of CLAMP-occupied sites throughout the genome before becoming enriched at its strongest X-linked target sites (CES). Synergistic interactions between CLAMP and MSLc (Albig et al., 2019; Soruco et al., 2013) are likely to enhance the occupancy of both factors at CES. We hypothesize that additional factors promote the specific interaction between MSLc and CES, including direct binding of MSL2 to DNA mediated by DNA shape (Albig et al. 2019), specific three-dimensional conformation surrounding CES (Ramírez et al. 2015), and 1.688 repetitive elements (Joshi and Meller 2017). In the future, it will be possible to use the meiotic drive system to define the contribution of each of these factors to MSLc targeting in a developmental context.

## ACKNOWLEDGEMENTS

We thank C. Staber and Bloomington Stock Center for the SD stocks and S. Blythe for the PCNA-eGFP stock. We thank Mitzi Kuroda for the goat anti-MSL3 serum. We thank the Hart lab at Brown University and the Dunn lab, formerly at Brown, now at Yale University, for use of their stereomicroscopes and S. Siebert for assistance taking embryo image in Figure 1C. We thank Dr. Marcela Soruco, Dr. Mattew Booker, Dr. Guray Kuzu, and Dr. Michael Tolstorukov for their important contributions. This work was supported by F32GM109663 and K99HD092625 to L.E.R., R35GM126994 to E.N.L., and an HHMI Gilliam fellowship and an NSF GRFP to W.T.J.

## EXPERIMENTAL PROCEDURES

### Fly stocks and crosses

Meiotic drive fly stocks (both gifts from Cynthia Staber), +; SD72/CyO and 19-3, yw, Rsp[s]-B[s]/Dp(2:y)CB25-4, y+, Rsp[s]B[s]; SPSD/CyO (Bloomington BSC64332) are published in (Rieder et al., 2017). To obtain female embryos, we mated +; SD72/CyO females to 19-3, yw, Rsp[s]-B[s]/Dp(2:y)CB25-4, y+, Rsp[s]B[s]; SPSD/CyO males to obtain +/Dp(2:y) CB25-4, y+, Rsp[s]B[s]; SPSD/SD72 males. To obtain male embryos, we mated +; SD72/CyO males to 19-3, yw, Rsp[s]-B[s]/Dp(2:y)CB25-4, y+, Rsp[s]B[s]; SPSD/CyO females to obtain 19-3, yw, Rsp[s]-B[s]/Y; SPSD/SD72 males. We mated males of both genotypes to yw; attP2{PCNA-EGFP} virgin females (Blythe and Wieschaus, 2016).

### Meiotic drive validation

*K* is the proportion of SD-bearing progeny compared to total progeny (Ganetzky, 1977; Gell and Reenan, 2013). In this case, *k* is expressed as the number of non-*Rsp*-bearing progeny, or the number of the *expected* sex, compared to total progeny (Figure 1B). We conducted *k-*tests at 24°C based on (Gell and Reenan, 2013) at by crossing either +/Dp(2:y) CB25-4, y+, Rsp[s]B[s]; SPSD/SD72 males or 19-3, yw, Rsp[s]-B[s]/Y; SPSD/SD72 males to yw; attP2{PCNA-EGFP} virgin females.

### Embryo fixation and sorting

We collected embryos on apple juice plates with yeast paste at 24°C. We performed 0-4hr timed lays and fixed embryos according to Blythe and Wieschaus (Blythe and Wieschaus, 2015). We then hand-sorted embryos using a Zeiss Discovery.V8 microscope under GFP excitation using an X-CITE 120Q stereo light source. We pooled 200-351 embryos (NC 11-14, **Figure 1D**). For 2-4hr embryos, we used approximately 100 mixed stage embryos. We froze embryos at -80°C until further processing.

### Chromatin immunoprecipitation (ChIP)-sequencing

We performed ChIP as in (Blythe and Wieschaus, 2015) using 2 μL of rabbit anti-CLAMP antibody, 2 μL of polyclonal rabbit IgG (Millipore-Sigma, 12-370), or 4 μL of a 1:10 dilution of goat anti-MSL3 serum (gift from Mitzi Kuroda) per sample. We synthesized libraries using the NEBNext ChIP-seq kit (New England Biosystems) and sequenced libraries on an Illumina HiSeq 2500 in 2×100-bp or 2×150-bp mode. ChIP-seq data is deposited at NCBI GEO and the accession number is pending.

### Datasets

Female embryo CLAMP ChIP-seq data are deposited in NCBI GEO under accession number GSE119448 (Rieder et al. 2017). Cell culture CLAMP ChIP-seq data are deposited in NCBI GEO under accession number GSE39271 (Soruco et al. 2013). We lifted CES locations (Soruco et al. 2013; Alekseyenko et al. 2008) over to dm6 using the UCSC liftOver tool.

### Chip-seq analysis

We mapped sequencing reads to release 6 of the *Drosophila melanogaster* genome (dm6) using Bowtie2 with default parameters. We identified reads with a MAPQ < 30 and removed PCR duplicate reads using Picard MarkDuplicates (version 2.9.2) using SAMtools (version 1.9). We used MACS2 (version 2.1.1) to identify peaks with the following parameters: --nomodel -B –SPMR –keep-dup all -g dm. We used input for peak calling in all samples with the exception of male 2-4 hour time point. We used a narrow peak calling with a q cutoff of 0.01 for CLAMP ChIPs and broad peak calling with a q cutoff of 0.05 for MSL3 and H4K16ac ChIPs; we also used --extsize of 147 for H4K16ac ChIPs. We used MACS2 to generate fold enrichment tracks for each ChIP. We generated average profiles using deepTools (version 3.1.0). We calculated distances to the nearest CES using bedtools (version 2.27.1). We determined peak overlaps using Intervene (Khan and Mathelier 2017).

---

**Figure S1.**
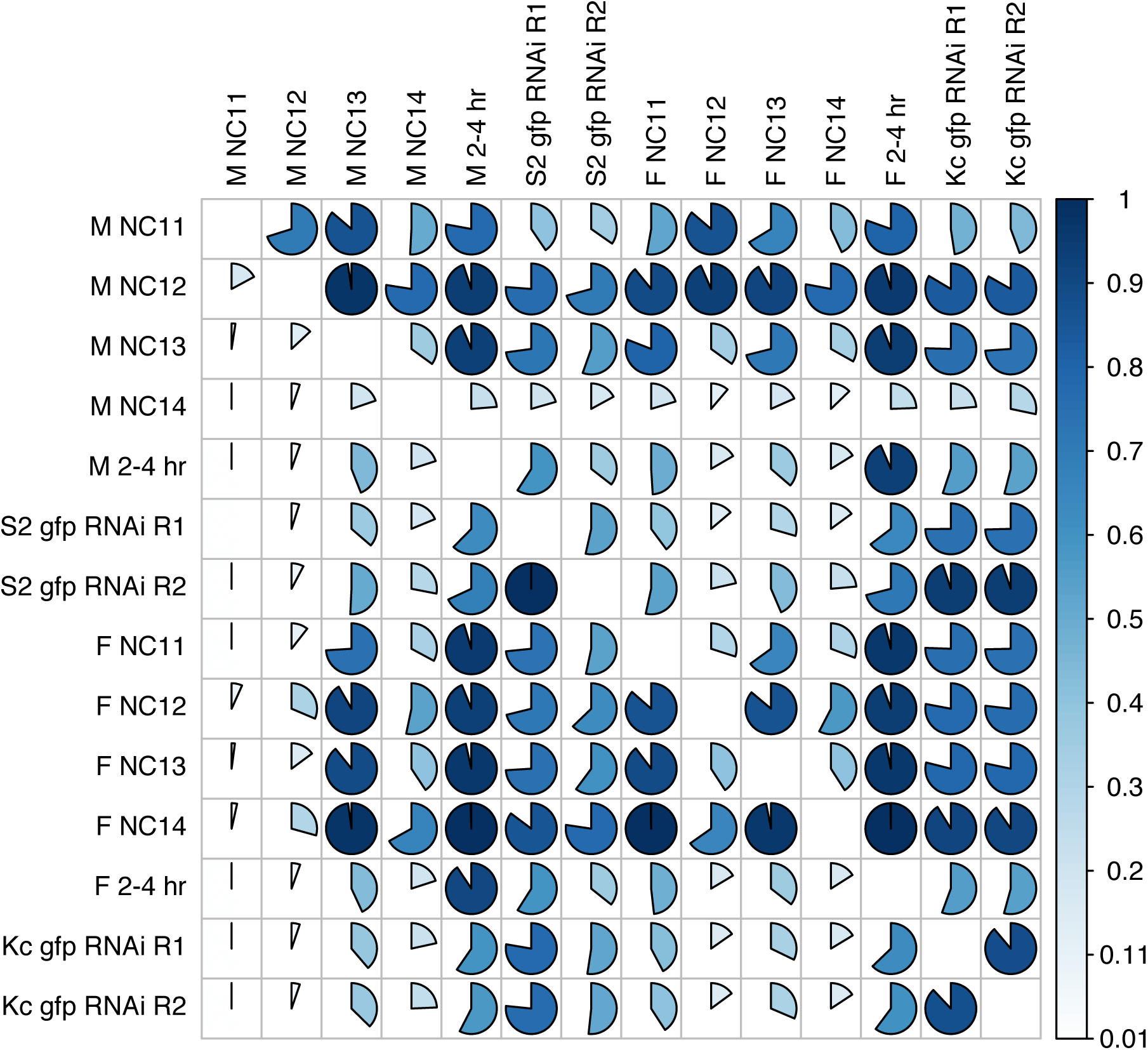
Percent overlap of CLAMP ChIP-seq peaks between all samples from the current study and previous cell culture CLAMP ChIP-seq samples. Cell culture (S2, male; Kc, female) CLAMP ChIP-seq data from Soruco et al. 2013, NCBI accession number GSE39271.

**Figure S2.**
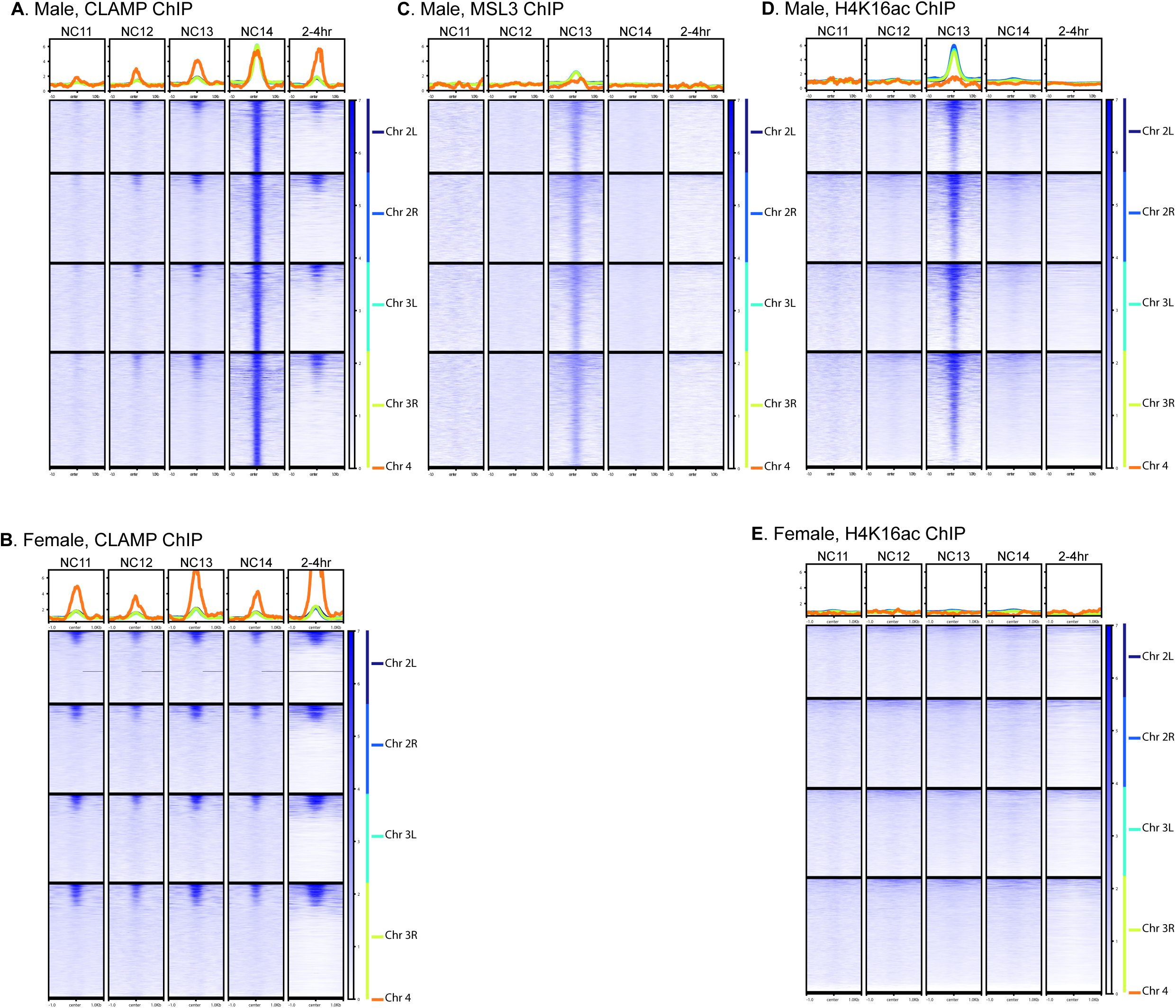
Staged, sexed ChIP-seq heat maps at autosomal CLAMP sites. Data are mapped over autosomal CLAMP sites and broken into chromosomes: 2L, dark blue; 2R, light blue; 3L, teal; 3R, light green; 4, orange. (A) CLAMP ChIP-seq from male embryos. (B) CLAMP ChIP-seq from female embryos. (C) MSL3 ChIP-seq from male embryos. (D) H4K16ac ChIP-seq from male embryos. (E) H4K16ac ChIP-seq from female embryos.

**Figure S3.**
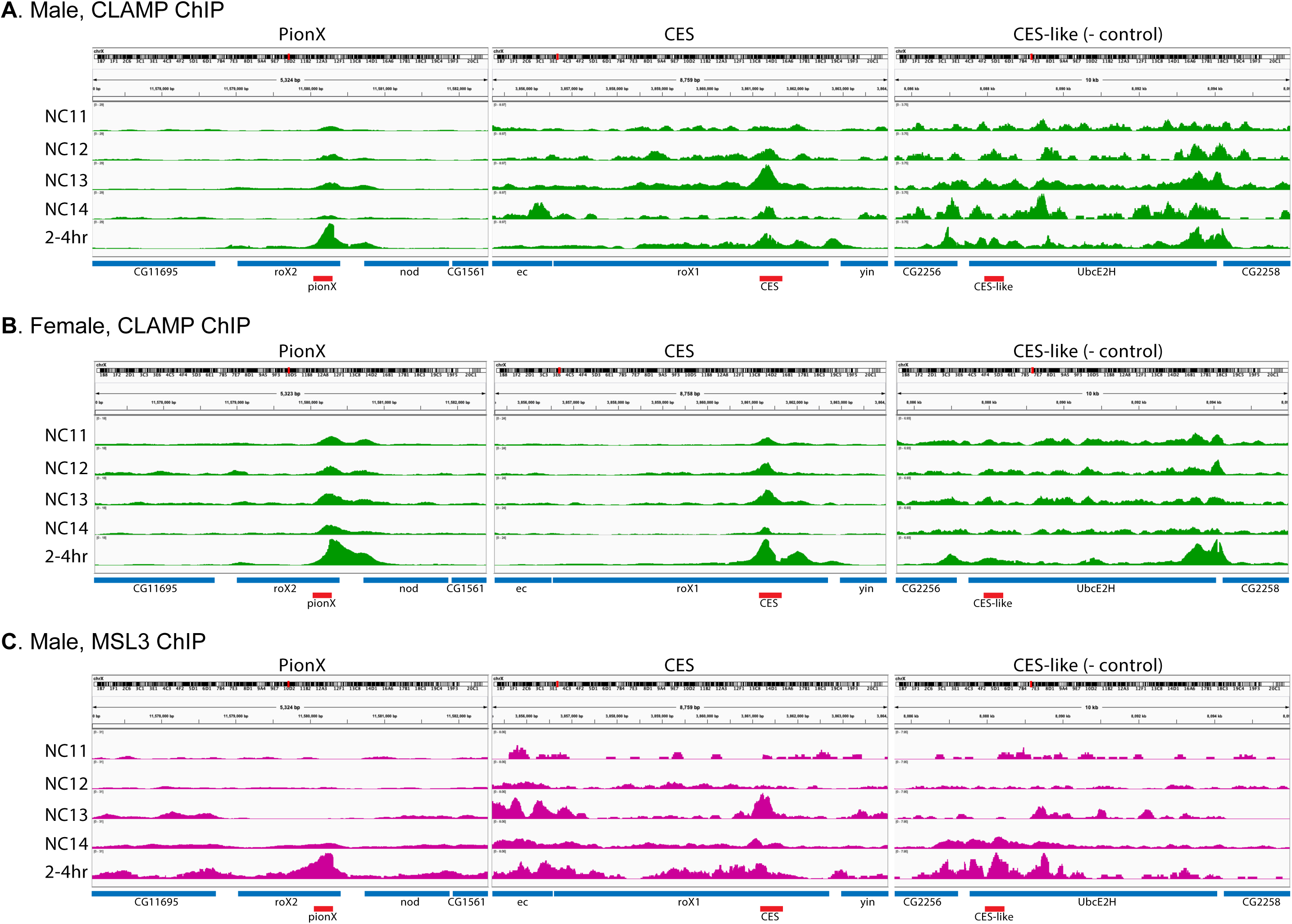
CLAMP (A and B) and MSL3 (C) ChIP-seq profiles over individual loci representative of classes in **Figure 2**, including *roX2* (PionX; Villa et al. 2016), *roX1* (CES; Soruco et al. 2013, Alekseyenko et al. 2008), and *UbcE2H* (CES-like; Soruco et al. 2013, Alekseyenko et al. 2008). Nearby genes are represented in blue and PionX, CES, and CES-like sites in red. Profiles generated in Integrative Genomics Viewer. Data are group autoscaled.

